# Atypical landscape of transposable elements in the large genome of *Aedes aegypti*

**DOI:** 10.1101/2024.02.07.579293

**Authors:** Josquin Daron, Alexander Bergman, Hélène Lopez-Maestre, Louis Lambrechts

**Author notes:** Corresponding author*: Josquin Daron.

## Abstract

Transposable elements (TEs) contribute significantly to variation in genome size among eukaryotic species, but the factors influencing TE accumulation and diversification are only partially understood. Most of our current knowledge about TE organization, dynamics and evolution derives from investigations in model organisms with a relatively small genome size such as *Drosophila melanogaster* or *Arabidopsis thaliana*. Whether the observed patterns hold true in larger genomes remains to be determined. The Diptera order is an ideal taxon to address this question, because it includes a forty-year model of TE biology (*D. melanogaster*) as well as mosquito species with significantly larger genomes. Here, we use a comparative genomics approach to characterize the genomic forces that have shaped the TE content of the *Aedes aegypti* genome (1.3 Gb) relative to the *Anopheles coluzzii* genome (300 Mb) and the *D. melanogaster* genome (180 Mb). Leveraging a newly developed high-quality TE library for *Ae. aegypti*, our results reveal a contrasted pattern of TE organization in *Ae. aegypti* compared to *An. coluzzii* and *D. melanogaster*. Our analyses suggest that the substantial TE fraction observed in the *Ae. aegypti* genome reflect both a high rate of TE transposition and a low rate of TE elimination. Together, our results indicate that TE organization and evolutionary dynamics in the large genome of *Ae. aegypti* are distinct from those of other dipterans with smaller genomes.

## Introduction

Transposable elements (TEs) are ubiquitous components of genomes found across all eukaryotic species except some *Plasmodium* parasites (reviewed in Wells & Feschotte, 2020). They consist of two major classes: retrotransposons (class I), which transpose via reverse transcription of an RNA intermediate, and DNA transposons (class II), which encompass all other types of TEs (Wicker *et al*., 2007). TEs are generally devoid of a function that would allow them to be maintained by natural selection, instead, their strategy relies on massive self-amplification from one genomic location to another (Le Rouzic & Capy, 2005; Hua-Van *et al*., 2011). This behavior is inherently mutagenic and generates a vast spectrum of genetic polymorphisms that can result in contrasting outcomes (Catlin & Josephs, 2022). While the mutagenic activity of TEs typically results in deleterious or neutral insertions, some TEs can be beneficial, playing a crucial role in adaptive evolution. The iconic example of the peppered moth illustrates how a TE’s influence on cortex gene expression drove adaptive changes in response to environmental factors during the Industrial Revolution (Hof *et al*., 2016). Accumulation of TEs in the genome represents a source of raw genetic material, causing a wide range of modifications in gene expression and function, such as the modeling of new regulatory networks and the creation of new genes (Feschotte, 2008; Cosby *et al*., 2021). Thus, TEs are considered one of the major drivers of genome evolution.

TE landscapes vary dramatically across species (Wells & Feschotte, 2020). Some genomes harbor only a few TEs, while others contain a broad and diverse repertoire of TEs. Since TE insertions are mostly deleterious, early work from Lynch and Conery hypothesized that TE content and genome size may be directly related to demographic constraints (Lynch & Conery, 2003). When the effective population size is small, genetic drift can cause large changes in allele frequencies that counteract the effect of purifying selection. Consequently, deleterious TE insertions that would be eliminated by purifying selection in larger populations can reach fixation in small populations. This hypothesis has been supported by empirical work showing that TEs seem to thrive in small populations (Lockton *et al*., 2008a; Stritt *et al*., 2018), especially in invasive populations where the efficiency of purifying selection is reduced due to the strong genetic bottlenecks imposed by founder events (Schrader *et al*., 2014; Mérel *et al*., 2021). However, this logic fails to explain differences in TE abundance and TE diversity observed between species with comparable effective population sizes, such as those within the same taxonomic order (Whitney *et al*., 2011; Lynch, 2011; Kapusta *et al*., 2017; Pasquesi *et al*., 2018). For instance, while several genera in the Diptera order have a compact genome, such as *Drosophila* or *Anopheles* (*D. melanogaster*, ∼0.18 Gb, dos Santos *et al*., 2015; *An. coluzzii*, ∼0.28 Gb, Vargas-Chavez *et al*., 2022), the *Aedes* genus stands out with genomes exceeding 1.0 Gb in size (*Ae. aegypti*, ∼1.3 Gb, Matthews *et al*., 2018). Yet the effective population size estimates of these three species typically range from 10^6^ for natural populations of *Ae. aegypti* (Rose *et al*., 2023) and *D. melanogaster* (Duchen *et al*., 2013; Sprengelmeyer *et al*., 2020) to 10^7^ for highly diverse *An. coluzzii* populations in West Africa (Ag1000G Consortium, 2017). The large variation in genome size despite a similar effective population size indicates that additional forces are at play, such as the rate of transposition and the rate at which TE sequences are eroded or deleted (Bourgeois & Boissinot, 2019). During the evolutionary history of a species, TE families escape silencing and experience bursts of amplification followed by rapid silencing (Daron *et al*., 2014; Kofler, 2019). The timing and intensity of such bursts are TE family-specific, which explains why TE families can vary in size from a single member to thousands of members (Wicker *et al*., 2018). Conversely, ectopic recombination can purge the genome from its TE content and prevent it from inflating under the action of TEs (Kapusta *et al*., 2017). Large genomes appear to lose DNA at a slower pace than smaller genomes, indicating that differences in the rate of DNA removal may play an important role in the variability of the TE content across species (Bensasson *et al*., 2001; Wicker & Keller, 2007; Hawkins *et al*., 2009; Sun *et al*., 2012).

Here, we examine the TE landscape in the large genome of *Ae. aegypti*, in comparison with that of *D. melanogaster* and *An. coluzzii*, two closely related dipteran species with smaller genomes. The mosquito *Ae. aegypti* is most famous as the primary vector of several human-pathogenic viruses including dengue, Zika, and yellow fever viruses. There is a long-standing interest in deciphering the *Ae. aegypti* genome because it is essential for advancing our understanding of vector biology and for developing effective strategies to mitigate the impact of mosquito-borne viruses on global public health (Severson *et al*., 2004). Although *Ae. aegypti* is one of the most tractable mosquito species for laboratory studies, progress in *Ae. aegypti* genomics has been hampered by the large genome size and high proportion of repeated sequences (Nene *et al*., 2007). Only recently has a high-quality reference genome been fully assembled and annotated using a combination of technologies (Matthews *et al*., 2018). Despite their overwhelming presence, however, little is known about the TE landscape of the *Ae. aegypti* genome. While some insights into the raw TE content were provided from studies whose main focus was horizontal TE transfer (Melo & Wallau, 2020) or TE regulation (Ma *et al*., 2021), basic knowledge is still lacking about the evolutionary dynamics of TEs and their organization in the *Ae. aegypti* genome. To begin addressing this knowledge gap, here we take advantage of a newly created high-quality TE library for *Ae. aegypti* and use a comparative genomics approach to characterize the genomic forces that have shaped the TE content in the genomes of *D. melanogaster, An. coluzzii* and *Ae. aegypti*. Our results reveal that the large TE content of *Ae. aegypti* likely results from accumulation of highly divergent TE copies. In contrast with the classic partitioning observed in small-genome organisms between TE-rich and TE-poor regions, TEs are more homogeneously distributed across the genomic landscape of *Ae. aegypti*.

## Results and Discussion

### Generating a unified and manually curated TE library for *Ae. aegypti*

To investigate the genomic landscape of TEs within the *Ae. aegypti* genome, we first generated an improved species-specific TE library using a combination of automated and manual methods (see Methods). Prior to our work, three main TE libraries were already available for *Ae. aegypti*. Two libraries were based on the *Ae. aegypti* reference genome, the TEfam library (Matthews *et al*., 2018) and a library generated by the Lau laboratory (Ma *et al*., 2021), which we refer to as “Lau” hereafter. A third TE library was derived from the genome sequence of the *Ae. aegypti* cell line Aag2 (Whitfield *et al*., 2017). We evaluated the performance of each of the three libraries to capture the TE repertoire of *Ae. aegypti* relative to a strictly *de novo* TE library created with the *RepeatModeler* program (Flynn *et al*., 2020). Although all four TE libraries differ widely in the total number of TE consensi (ranging from 1090 to 3462, Figure 1A), the genomic fraction they cover is similar between them (from 54% to 66%, Figure 1B). This indicates that although the TE libraries represent a consistent amount of TE sequences present in the genome, they have different levels of redundancy and/or fragmentation. For instance, the significantly higher numbers of TE consensi in the Aag2 and *de novo* libraries are likely due to the presence of incomplete consensi that remain to be concatenated.

**Figure 1:**
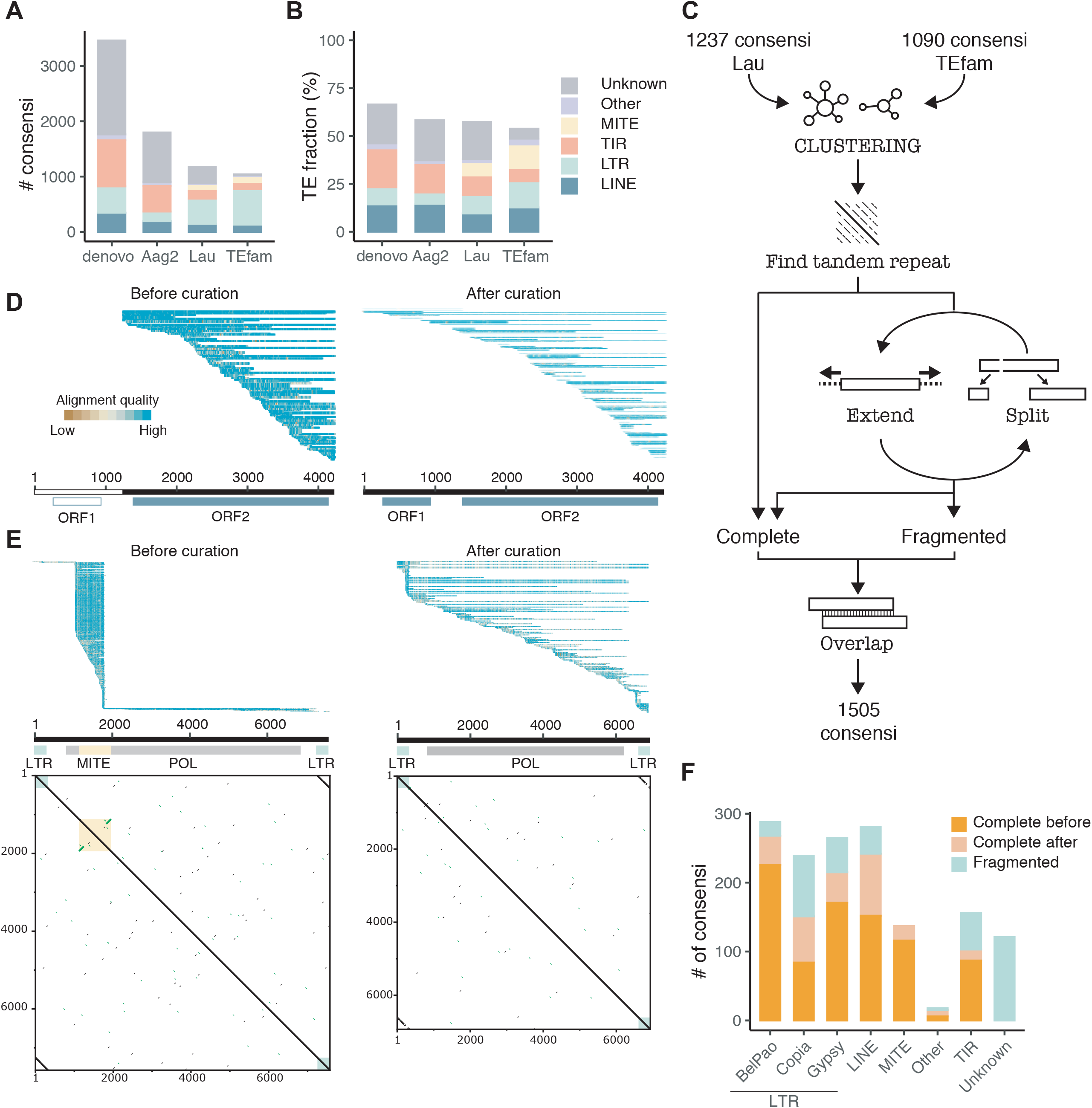
Creation of a high-quality TE library for *Ae. aegypti*. (A) Number of TE consensi (color-coded by superfamily) in the *de novo*, Aag2, Lau and TEfam libraries. (B) Fraction of the *Ae. aegypti* genome masked by each TE library. (C) Workflow of the automated and manual steps to generate a high-quality TE library. (D) Pairwise alignment of genomic TE copies on an example LINE retrotransposon consensus sequence before and after curation. Each sequence is represented by a single row (sorted by start position) where the color gradient indicates alignment quality over non-overlapping 10-b windows. (E) Example nested structure of a MITE within an LTR retrotransposon with the same display as panel D but with all-against-all dotplot visualization below the pairwise alignment. LTRs and TIRs are visualized by the short line parallel or perpendicular to the 1-to-1 line. (F) Barplot summarizing the improvement of consensus completeness in the final curated TE library.

To generate a unified TE library for *Ae. aegypti*, we first merged the TEfam and Lau libraries because they had the most complete set of TE consensi among the existing libraries. We then improved the quality of the TE consensi by clustering and manual curation through an *in house* workflow (Figure 1C), which resulted in a single TE library of 1505 consensi. At the heart of this workflow, a semi-automated curation procedure was used to recover a complete edge-to-edge TE structure of the fragmented consensi. Figure 1D and 1E illustrate two critical cases of such improvements. Figure 1D displays a truncated long interspersed nuclear element (LINE) consensus only containing the open reading frame 2 (ORF2), with the pairwise alignment of genomic TE copies exhibiting typical 5’ truncated pattern. We extended the 5’ end of the consensus using the *alignAndCallCons* program to recover complete edge-to-edge LINE structure. Figure 1E illustrates the presence of a miniature inverted-repeat TE (MITE) nested with a long terminal repeat (LTR) retrotransposon consensus. The presence of the MITE was revealed by dotplot visualization, and the stack of genomic copies mapped precisely to the MITE sequence. Numerous similar cases were observed, for which we carefully manually removed the MITE sequence from the consensus sequence. Together, this workflow allowed us to retrieve a complete edge-to-edge TE structure for 43% of the initial fragmented consensi (Figure 1F) with the most significant improvement for LTR and LINE retrotransposons. We note that the final TE library still includes 121 consensi with unknown classification. These consensi are repeated sequences without an obvious structural feature that would allow their assignment to a TE family. This is likely common in genomes with high TE content, where the nested structure of TEs (insertions within insertions) results in a high level of fragmentation.

### Inefficient purging mechanisms likely drive the accumulation of TEs in *Ae. aegypti*

To investigate the genomic forces underlying the large TE content of the *Ae. aegypti* genome, we compared the TE landscape of *Ae. aegypti* with that of two dipterans with smaller genomes, *An. coluzzii* and *D. melanogaster*, through the lens of a comparative genomics approach. These two species were chosen for the availability of their high-quality reference genomes and TE libraries. Collectively, interspersed TEs account for ∼60% of the *Ae. aegypti* genome sequence (Figure 2B), with LINEs representing 15.37%, followed by MITEs (13.10%), LTR retrotransposons (11.31%), and TIR elements (9.02%) (Figure 2C). As expected, the TE fraction in *Ae. aegypti* represents a three-to five-fold increase relative to *An. coluzzii* and *D. melanogaster*, respectively. The sharp contrast in TE content between *Ae. aegypti* and the small-genome dipterans is mostly driven by the higher abundance of LINE, LTR *copia*, TIR elements, and of non-autonomous MITEs. However, the genomic fraction of LTR *gypsy* elements is remarkably higher in *D. melanogaster* (5.34%) than in *Ae. aegypti* (3.51%). Together, these results add up to the accumulating empirical evidence that the evolutionary success of TE superfamilies is species-specific. The contemporary effective population sizes of the three species have been estimated to range from 10^6^ to 10^7^ (Rose *et al*., 2023; Ag1000G Consortium, 2017; Duchen *et al*., 2013; Sprengelmeyer *et al*., 2020). Thus, it is unlikely that the large discrepancy in TE content among the three species is explained by major differences in the current efficiency of purifying selection to remove deleterious mutations. We cannot rule out, however, that accumulation of TEs in several *Aedes* mosquito species results from a population bottleneck that preceded the radiation of the *Aedes* genus.

**Figure 2:**
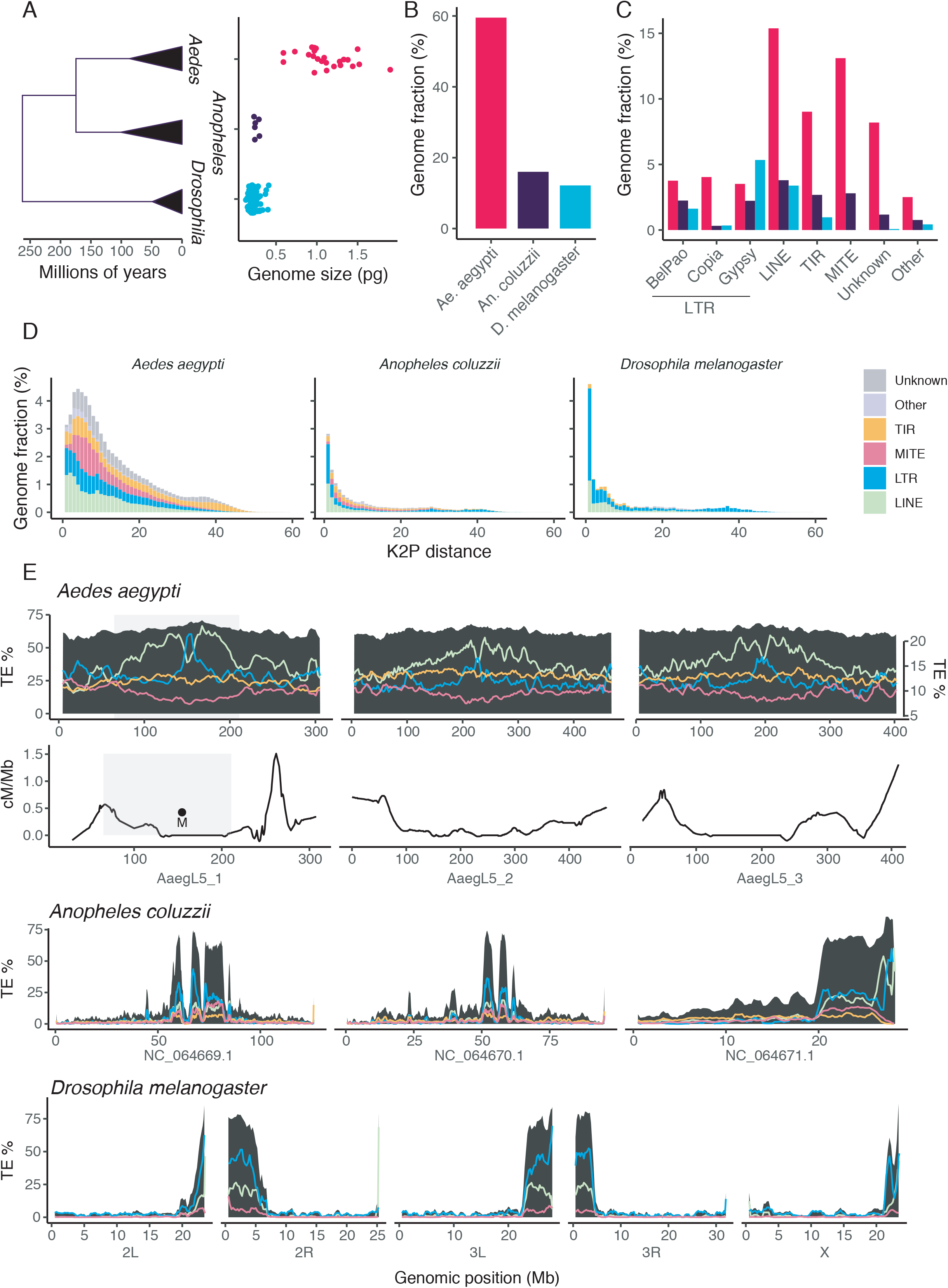
Comparative genomics of the TE landscape in *Ae. aegypti, An. coluzzii* and *D. melanogaster*. (A) Schematic phylogenetic relationships between the *Drosophila, Anopheles* and *Aedes* genera. Branch lengths in millions of years were obtained from Soghigian *et al*., 2023. Scatterplot showing the distribution of the haploid DNA contents (C-values, in picograms) collected form the Animal Genome Size Database (www.genomesize.com). (B) Total TE fraction of the *Ae. aegypti* (red), *An. coluzzii* (dark blue) and *D. melanogaster* (light blue) genomes. (C) TE fraction in the three genomes stratified by superfamilies. (D) Age structure of TEs (color-coded by superfamily) shown as the frequency distribution of copies as a function of their CpG-corrected Kimura divergence from the consensus (K2P, Kimura 2-parameter metric) in the three species. (E) Genomic density of TEs (black area) or TE superfamilies (lines color-coded according to the legend of panel D) in overlapping windows of 10% of the genome size with a step of 1%, shown as percentages along the chromosomes. For *Ae. aegypti*, the secondary y-axis denotes the genomic density of TE superfamilies. For *Ae. aegypti*, the meiotic recombination rate in cM/Mb is shown using data from Chen *et al*., 2022. The overlaid light grey shading indicates the 123-Mb region of reduced male recombination identified by Fontaine *et al*., 2017 around the male-determining (M) locus on chromosome 1.

To determine whether the higher TE content of *Ae. aegypti* reflects a higher rate of transposition and/or a reduced rate of TE elimination, we examined the evolutionary dynamics of TEs based on the age structure of TE copies (Kimura, 1980). We used within-subfamily divergence as a proxy for TE subfamily age, stratified by TE superfamily (Figure 2D). Interestingly, the higher TE content in *Ae. aegypti* is not associated with an excess of very recent TE copies, since the genomic fraction of young TEs (i.e., with a sequence identical to the consensus) is in the same range for *D. melanogaster, An. coluzzii* and *Ae. aegypti* (4.6%, 2.8% and 3.1%, respectively). In contrast, most of TE sequences in *Ae. aegypti* are highly divergent from the consensus. This pattern is relatively consistent across all TE superfamilies, although some variation exists. Together, these observations suggest that the higher TE content in *Ae. aegypti* is primarily due to a lower rate of TE purging after insertion. However, this interpretation assumes that TE insertion and elimination rates are constant over time. In future work, it will be interesting to model temporal variation of TE transposition and elimination rates and untangle their relative contributions (Bourgeois & Boissinot, 2019).

To compare the spatial distribution of TEs along the chromosomes in the three dipteran species, we examined variation in their local density in overlapping windows of 10% of the genome size with a step of 1% (Figure 2E). TE distribution in the genomes of *An. coluzzii* and *D. melanogaster* is highly clustered, with high TE densities in pericentromeric regions and low TE densities in distal areas of the chromosomes. For instance, TE density varies from 0.61% to 81.2% in *D. melanogaster*. This is in sharp contrast with TE distribution in the *Ae. aegypti* genome, which is remarkably even along the chromosomes, ranging from 52.9% to 67.8%. Within the overall TE distribution, each superfamily exhibits its own density pattern. TIR elements and their non-autonomous MITE versions follow the overall even distribution, whereas LINE and LTR retrotransposons reach higher densities in the pericentromeric region of the chromosomes. Increased accumulation of retrotransposons is particularly noticeable within the 123-Mb region of chromosome 1 that contains the sex-determining locus and displays reduced male recombination in *Ae. aegypti* (Fontaine *et al*., 2017; Chen *et al*., 2022). Remarkably, a localized inversion of TE accumulation between LTR and LINE retrotransposons occurs at the precise genomic location of the male-determining (M) locus on chromosome 1. The other two chromosomes display similar but less pronounced patterns in the low recombination regions around the centromeres. Our results are consistent with previous studies reporting that TEs tend to accumulate in genomic regions of low recombination and to be purged by purifying selection from regions with a higher rate of recombination (Sniegowski & Charlesworth, 1994; Barrón *et al*., 2014; Charlesworth & Campos, 2014).

Although TE transposition seems to be ongoing in the *Ae. aegypti* genome, the fraction of recently duplicated TE copies –lower than the fraction in *D. melanogaster*– cannot explain alone the disproportionately higher TE content. The widespread presence of highly divergent TE copies that have accumulated over time points to a low rate of TE elimination, possibly due to the relatively lower meiotic recombination rate of *Ae. aegypti* (∼0.3 cM/Mb) compared to *An. coluzzii* (∼0.8 cM/Mb) and *D. melanogaster* (∼1.6 cM/Mb) (Wilfert *et al*., 2007). Together, our analyses indicate that the high TE content of the *Ae. aegypti* genome results from both a sustained rate of new TE insertions and inefficient purifying selection. More generally, our results indicate that TE organization and evolutionary dynamics in the large genome of *Ae. aegypti* are distinct from those of other dipterans with smaller genomes.

## Methods

### TE library

We used a combination of automated and manual techniques to generate a high-quality TE library for the *Ae. aegypti* reference genome (Figure 1C). Briefly, TE consensi from TEfam (Matthews *et al*., 2018), and from the TE library generated by Ma *et al*., 2021 (referred to as the Lau library) were clustered using *CD-HIT-EST* (-c 0.8 -aS 0.8 -b 500) to generate a non-redundant set of consensi (Fu *et al*., 2012). Manual inspection with *dotter* (Sonnhammer & Durbin, 1995) of this non-redundant set revealed the presence of simple tandem repeats, which we masked using *Tandem Repeats Finder* (Benson, 1999). Simple tandem repeats representing over 50% of a consensus were excluded from the downstream analysis. Next, TE consensi were divided into two groups, complete or fragmented, based on the presence of structural features annotated in their sequences (LTR, TIR, protein-coding domains). LTR and TIR sequences were identified by manual inspection with *dotter*, while protein-coding domains were annotated by *blastx* (Altschul *et al*., 1990) against the *RepeatMasker* protein database (www.repeatmasker.org/RepeatProteinMask.html#database). LTR or TIR element consensi were considered as complete when they were flanked by a pair of LTRs/TIRs, and their internal sequence contained the relevant protein-coding domains. Complete LINE retrotransposon consensi were defined by the presence of two complete ORFs. To include the most complete consensi as possible in our library, we used the program *alignAndCallCons* (-d 40 -ma 20 -re 100 -fi -ht -p 5 -f 3) from the Dfam toolbox (Storer *et al*., 2021) to extend the edge of fragmented consensi until reaching a complete structure as previously defined. In parallel, manual inspection of *dotter* display and pairwise alignment of genomic copies aligned to the consensus sequence revealed the presence of several chimeric consensi derived from the juxtaposition of two different TEs. These chimeric consensi included MITEs nested within other consensi. We treated chimeric consensi in two ways: (i) MITEs were excised from the larger consensus, and (ii) juxtaposed TE sequences were split and fed back to the extension step to obtain an edge-to-edge TE structure. Finally, to remove any redundancy created by the numerous manipulations, we performed an all-versus-all *blastn* of the library to detect and remove overlapping consensi based on the 80-80-80 rule criterion (Wicker *et al*., 2007). We also generated a *de novo* TE library with *RepeatModeler* (Flynn *et al*., 2020) to serve as a reference. The newly created unified TE library for *Ae. aegypti* is available upon request.

### TE annotation and genomic analysis

We used the standard similarity search approach using *RepeatMasker* to annotate TEs in the reference genomes of *Ae. aegypti* (AaegL5.3, Matthews *et al*., 2018), *An. coluzzii* (acolN3, Vargas-Chavez *et al*., 2022) and *D. melanogaster* (r6.46, dos Santos *et al*., 2015), using a TE library specifically dedicated to each genome. As an extra round of polishing, we used the *CLARITE* program (Daron *et al*., 2014) to improve TE modeling and reconstruct their nested structure. CpG-corrected Kimura divergence from the consensus was collected from the output of *RepeatMasker*. Estimates of the TE density along the genomes were computed using *bedtools intersect* using sliding windows of 10% of the genome size with a step of 1%.

## Acknowledgements

This work was supported by the French Government’s Investissement d’Avenir program, Laboratoire d’Excellence Integrative Biology of Emerging Infectious Diseases (grant ANR-10-LABX-62-IBEID) and the Agence Nationale de la Recherche (grant ANR-18-CE35-0003-01). A. Bergman was supported by a stipend from the Pasteur - Paris University (PPU) International PhD Program. We acknowledge the HPC Core Facility at Institut Pasteur for providing HPC resources that have contributed to the research results reported within this paper. The funders had no role in study design, data collection and analysis, decision to publish, or preparation of the manuscript.

## References

Ag1000G Consortium (2017) Genetic diversity of the African malaria vector Anopheles gambiae. Nature 552: 96–100

Altschul SF, Gish W, Miller W, Myers EW & Lipman DJ (1990) Basic local alignment search tool. J Mol Biol 215: 403–410

Barrón MG, Fiston-Lavier A-S, Petrov DA & González J (2014) Population Genomics of Transposable Elements in Drosophila. Annu Rev Genet 48: 561–581

Bensasson D, Petrov DA, Zhang D-X, Hartl DL & Hewitt GM (2001) Genomic Gigantism: DNA Loss Is Slow in Mountain Grasshoppers. Mol Biol Evol 18: 246–253

Benson G (1999) Tandem repeats finder: a program to analyze DNA sequences. Nucleic Acids Res 27: 573–580

Bourgeois Y & Boissinot S (2019) On the Population Dynamics of Junk: A Review on the Population Genomics of Transposable Elements. Genes 10: 419

Catlin NS & Josephs EB (2022) The important contribution of transposable elements to phenotypic variation and evolution. Curr Opin Plant Biol 65: 102140

Charlesworth B & Campos JL (2014) The Relations Between Recombination Rate and Patterns of Molecular Variation and Evolution in Drosophila. Annu Rev Genet 48: 383–403

Chen C, Compton A, Nikolouli K, Wang A, Aryan A, Sharma A, Qi Y, Dellinger C, Hempel M, Potters M, et al. (2022) Marker-assisted mapping enables forward genetic analysis in Aedes aegypti, an arboviral vector with vast recombination deserts. Genetics 222: iyac140

Cosby RL, Judd J, Zhang R, Zhong A, Garry N, Pritham EJ & Feschotte C (2021) Recurrent evolution of vertebrate transcription factors by transposase capture. Science 371: eabc6405

Daron J, Glover N, Pingault L, Theil S, Jamilloux V, Paux E, Barbe V, Mangenot S, Alberti A, Wincker P, et al. (2014) Organization and evolution of transposable elements along the bread wheat chromosome 3B. Genome Biol 15: 546

dos Santos G, Schroeder AJ, Goodman JL, Strelets VB, Crosby MA, Thurmond J, Emmert DB, Gelbart WM, & the FlyBase Consortium (2015) FlyBase: introduction of the Drosophila melanogaster Release 6 reference genome assembly and large-scale migration of genome annotations. Nucleic Acids Res 43: D690–D697

Duchen P, Živković D, Hutter S, Stephan W & Laurent S (2013) Demographic Inference Reveals African and European Admixture in the North American Drosophila melanogaster Population. Genetics 193: 291–301

Feschotte C (2008) Transposable elements and the evolution of regulatory networks. Nat Rev Genet 9: 397–405

Flynn JM, Hubley R, Goubert C, Rosen J, Clark AG, Feschotte C & Smit AF (2020) RepeatModeler2 for automated genomic discovery of transposable element families. Proc Natl Acad Sci 117: 9451–9457

Fontaine A, Filipović I, Fansiri T, Hoffmann AA, Cheng C, Kirkpatrick M, Rašić G & Lambrechts L (2017) Extensive Genetic Differentiation between Homomorphic Sex Chromosomes in the Mosquito Vector, Aedes aegypti. Genome Biol Evol 9: 2322–2335

Fu L, Niu B, Zhu Z, Wu S & Li W (2012) CD-HIT: accelerated for clustering the next-generation sequencing data. Bioinformatics 28: 3150–3152

Hawkins JS, Proulx SR, Rapp RA & Wendel JF (2009) Rapid DNA loss as a counterbalance to genome expansion through retrotransposon proliferation in plants. Proc Natl Acad Sci 106: 17811–17816

Hof AE van’t, Campagne P, Rigden DJ, Yung CJ, Lingley J, Quail MA, Hall N, Darby AC & Saccheri IJ (2016) The industrial melanism mutation in British peppered moths is a transposable element. Nature 534: 102–105

Hua-Van A, Le Rouzic A, Boutin TS, Filée J & Capy P (2011) The struggle for life of the genome’s selfish architects. Biol Direct 6: 19

Kapusta A, Suh A & Feschotte C (2017) Dynamics of genome size evolution in birds and mammals. Proc Natl Acad Sci 114: E1460–E1469

Kimura M (1980) A simple method for estimating evolutionary rates of base substitutions through comparative studies of nucleotide sequences. J Mol Evol 16: 111–120

Kofler R (2019) Dynamics of Transposable Element Invasions with piRNA Clusters. Mol Biol Evol 36: 1457–1472

Le Rouzic A & Capy P (2005) The First Steps of Transposable Elements Invasion: Parasitic Strategy vs. Genetic Drift. Genetics 169: 1033–1043

Lockton S, Ross-Ibarra J & Gaut BS (2008a) Demography and weak selection drive patterns of transposable element diversity in natural populations of Arabidopsis lyrata. Proc Natl Acad Sci 105: 13965–13970

Lockton S, Ross-Ibarra J & Gaut BS (2008b) Demography and weak selection drive patterns of transposable element diversity in natural populations of Arabidopsis lyrata. Proc Natl Acad Sci 105: 13965–13970

Lynch M (2011) Statistical Inference on the Mechanisms of Genome Evolution. PLOS Genet 7: e1001389

Lynch M & Conery JS (2003) The Origins of Genome Complexity. Science 302: 1401–1404

Ma Q, Srivastav SP, Gamez S, Dayama G, Feitosa-Suntheimer F, Patterson EI, Johnson RM, Matson EM, Gold AS, Brackney DE, et al. (2021) A mosquito small RNA genomics resource reveals dynamic evolution and host responses to viruses and transposons. Genome Res 31: 512–528

Matthews BJ, Dudchenko O, Kingan SB, Koren S, Antoshechkin I, Crawford JE, Glassford WJ, Herre M, Redmond SN, Rose NH, et al. (2018) Improved reference genome of Aedes aegypti informs arbovirus vector control. Nature 563: 501–507

Melo ES de & Wallau GL (2020) Mosquito genomes are frequently invaded by transposable elements through horizontal transfer. PLOS Genet 16: e1008946

Mérel V, Gibert P, Buch I, Rodriguez Rada V, Estoup A, Gautier M, Fablet M, Boulesteix M & Vieira C (2021) The Worldwide Invasion of Drosophila suzukii Is Accompanied by a Large Increase of Transposable Element Load and a Small Number of Putatively Adaptive Insertions. Mol Biol Evol 38: 4252–4267

Nene V, Wortman JR, Lawson D, Haas B, Kodira C, Tu Z (Jake), Loftus B, Xi Z, Megy K, Grabherr M, et al. (2007) Genome Sequence of Aedes aegypti, a Major Arbovirus Vector. Science 316: 1718–1723

Pasquesi GIM, Adams RH, Card DC, Schield DR, Corbin AB, Perry BW, Reyes-Velasco J, Ruggiero RP, Vandewege MW, Shortt JA, et al. (2018) Squamate reptiles challenge paradigms of genomic repeat element evolution set by birds and mammals. Nat Commun 9: 2774

Rose NH, Badolo A, Sylla M, Akorli J, Otoo S, Gloria-Soria A, Powell JR, White BJ, Crawford JE & McBride CS (2023) Dating the origin and spread of specialization on human hosts in Aedes aegypti mosquitoes. eLife 12: e83524

Schrader L, Kim JW, Ence D, Zimin A, Klein A, Wyschetzki K, Weichselgartner T, Kemena C, Stökl J, Schultner E, et al. (2014) Transposable element islands facilitate adaptation to novel environments in an invasive species. Nat Commun 5: 5495

Severson DW, Knudson DL, Soares MB & Loftus BJ (2004) Aedes aegypti genomics. Insect Biochem Mol Biol 34: 715–721

Sniegowski PD & Charlesworth B (1994) Transposable element numbers in cosmopolitan inversions from a natural population of Drosophila melanogaster. Genetics 137: 815–827

Soghigian J, Sither C, Justi SA, Morinaga G, Cassel BK, Vitek CJ, Livdahl T, Xia S, Gloria-Soria A, Powell JR, et al. (2023) Phylogenomics reveals the history of host use in mosquitoes. Nat Commun 14: 6252

Sonnhammer ELL & Durbin R (1995) A dot-matrix program with dynamic threshold control suited for genomic DNA and protein sequence analysis. Gene 167: GC1–GC10

Sprengelmeyer QD, Mansourian S, Lange JD, Matute DR, Cooper BS, Jirle EV, Stensmyr MC & Pool JE (2020) Recurrent Collection of Drosophila melanogaster from Wild African Environments and Genomic Insights into Species History. Mol Biol Evol 37: 627–638

Storer JM, Hubley R, Rosen J & Smit AFA (2021) Curation Guidelines for de novo Generated Transposable Element Families. Curr Protoc 1: e154

Stritt C, Gordon SP, Wicker T, Vogel JP & Roulin AC (2018) Recent Activity in Expanding Populations and Purifying Selection Have Shaped Transposable Element Landscapes across Natural Accessions of the Mediterranean Grass Brachypodium distachyon. Genome Biol Evol 10: 304–318

Sun C, López Arriaza JR & Mueller RL (2012) Slow DNA Loss in the Gigantic Genomes of Salamanders. Genome Biol Evol 4: 1340–1348

Vargas-Chavez C, Pendy NML, Nsango SE, Aguilera L, Ayala D & González J (2022) Transposable element variants and their potential adaptive impact in urban populations of the malaria vector Anopheles coluzzii. Genome Res 32: 189–202

Wells JN & Feschotte C (2020) A Field Guide to Eukaryotic Transposable Elements. Annu Rev Genet 54: 539–561

Whitfield ZJ, Dolan PT, Kunitomi M, Tassetto M, Seetin MG, Oh S, Heiner C, Paxinos E & Andino R (2017) The diversity, structure and function of heritable adaptive immunity sequences in the Aedes aegypti genome. Curr Biol CB 27: 3511-3519.e7

Whitney KD, Boussau B, Baack EJ & Jr TG (2011) Drift and Genome Complexity Revisited. PLOS Genet 7: e1002092

Wicker T, Gundlach H, Spannagl M, Uauy C, Borrill P, Ramírez-González RH, De Oliveira R, Mayer KFX, Paux E, Choulet F, et al. (2018) Impact of transposable elements on genome structure and evolution in bread wheat. Genome Biol 19: 103

Wicker T & Keller B (2007) Genome-wide comparative analysis of copia retrotransposons in Triticeae, rice, and Arabidopsis reveals conserved ancient evolutionary lineages and distinct dynamics of individual copia families. Genome Res 17: 1072–1081

Wicker T, Sabot F, Hua-Van A, Bennetzen JL, Capy P, Chalhoub B, Flavell A, Leroy P, Morgante M, Panaud O, et al. (2007) A unified classification system for eukaryotic transposable elements. Nat Rev Genet 8: 973–982

Wilfert L, Gadau J & Schmid-Hempel P (2007) Variation in genomic recombination rates among animal taxa and the case of social insects. Heredity 98: 189–197

